# SARS-CoV-2 selectively induces the expression of unproductive splicing isoforms of interferon, class I MHC and splicing machinery genes

**DOI:** 10.1101/2023.04.12.536671

**Authors:** Thomaz Lüscher Dias, Izabela Mamede Costa Andrade da Conceição, Nayara Evelin de Toledo, Lúcio Rezende Queiroz, Ícaro Castro, Rafael Polidoro Alves Barbosa, Luiz Eduardo Del-Bem, Helder Nakaya, Glória Regina Franco

**Author notes:** These authors contributed equally to this work (Thomaz Lüscher Dias, Izabela Mamede Costa Andrade da Conceição).

## Abstract

Splicing is a highly conserved, intricate mechanism intimately linked to transcription elongation, serving as a pivotal regulator of gene expression. Alternative splicing may generate specific transcripts incapable of undergoing translation into proteins, designated as unproductive. A plethora of respiratory viruses, including Severe Acute Respiratory Syndrome Coronavirus 2 (SARS-CoV-2), strategically manipulate the host’s splicing machinery to circumvent antiviral responses. During the infection, SARS-CoV-2 effectively suppresses interferon (IFN) expression, leading to B cell and CD8+ T cell leukopenia, while simultaneously increasing the presence of macrophages and neutrophils in patients with severe COVID-19. In this study, we integrated publicly available omics datasets to systematically analyze transcripts at the isoform level and delineate the nascent-peptide translatome landscapes of SARS-CoV-2-infected human cells. Our findings reveal a hitherto uncharacterized mechanism whereby SARS-CoV-2 infection induces the predominant expression of unproductive splicing isoforms in key IFN signaling genes, interferon-stimulated genes (ISGs), class I MHC genes, and splicing machinery genes, including IRF7, OAS3, HLA-B, and HNRNPH1. In stark contrast, cytokine and chemokine genes, such as IL6, CXCL8, and TNF, predominantly express productive (protein-coding) splicing isoforms in response to SARS-CoV-2 infection. We postulate that SARS-CoV-2 employs a previously unreported tactic of exploiting the host splicing machinery to bolster viral replication and subvert the immune response by selectively upregulating unproductive splicing isoforms from antigen presentation and antiviral response genes. Our study sheds new light on the molecular interplay between SARS-CoV-2 and the host immune system, offering a foundation for the development of novel therapeutic strategies to combat COVID-19.

## Introduction

Severe acute respiratory syndrome coronavirus 2 (SARS-CoV-2) is the pathogen that causes the coronavirus disease 2019 (COVID-19), which to date has killed over 6.7 million people worldwide. SARS-CoV-2 infects the upper respiratory tract and lungs and triggers a systemic inflammatory response that can lead to respiratory failure and death in severe cases (Carvalho et al. 2021). Unlike other coronaviruses and viruses known to interfere with host splicing machinery, such as Herpes Simplex Virus type 1, Epstein-Barr virus, and Yellow Fever virus, SARS-CoV-2 uniquely blocks interferon (IFN) expression and modulates immune cell populations (Blanco-Melo et al. 2020; Lokugamage et al. 2020; Miorin et al. 2020; Liao et al. 2020; Sandri-Goldin 2008; Verma et al. 2010; Bronzoni et al. 2011). Given the demonstrated ability of SARS-CoV-2 to interfere with host splicing machinery (Banerjee et al. 2020; Zaffagni et al. 2022), understanding the consequences of perturbations in splicing and unproductive splicing isoforms for the host’s antiviral response is crucial. Elucidating these mechanisms may reveal novel therapeutic targets and help develop strategies to counteract the virus’s immune evasion tactics.

In this study, we particularly focus on unproductive splicing isoforms, which are transcripts that do not code for functional proteins due to alterations in the splicing process. Unproductive splicing isoforms can result from various splicing events, such as intron retention, premature translation-termination codons (PTCs), or alternative splice site selection that disrupts the open reading frame (ORF). These unproductive isoforms can be targeted for nonsense-mediated decay (NMD) or remain untranslated in the nucleus, ultimately reducing protein production.

In light of these observations, our study aims to investigate the transcriptome-wide, isoform-level consequences of SARS-CoV-2-induced disruption in splicing, particularly focusing on the expression of unproductive splicing isoforms of key antiviral genes. We hypothesize that SARS-CoV-2 selectively manipulates the host splicing machinery to promote viral replication and subvert the immune response. We utilized cutting-edge bioinformatics approaches and publicly available omics datasets to gain a comprehensive understanding of the consequences of SARS-CoV-2 infection on host splicing. Our integrative analysis uniquely combines transcriptomic and nascent-peptide translatome data, enabling a more detailed characterization of the infected host cell landscape.

We found that SARS-CoV-2 infection induces a predominant expression of unproductive splicing isoforms of key IFN signaling genes and ISGs, class I MHC genes, and splicing machinery genes, such as IRF7, OAS3, HLA-B, and HNRNPH1. In contrast, this is not detected for cytokine and chemokine genes, such as IL6, CXCL8, and TNF, which preferentially express productive (protein-coding) splicing isoforms in response to SARS-CoV-2. We further found that several genes overexpressing unproductive splicing isoforms upon SARS-CoV-2 infection also show reduced protein synthesis in infected cells, including RBM5 and SRSF5. Finally, we demonstrated that many proteins and mRNAs affected by the SARS-CoV-2 disruption of splicing described here are also direct targets of viral proteins according to two distinct protein-protein interaction networks.

Our findings have significant implications for the development of therapeutic strategies against COVID-19. By uncovering the molecular interplay between SARS-CoV-2 and host splicing machinery, we provide novel insights into potential targets for antiviral drugs and immunomodulatory interventions. These discoveries may guide the design of therapies aimed at restoring normal splicing during viral infections.

## Results & Discussion

### SARS-CoV-2 infection increases the abundance of unproductive splicing isoforms

We first analyzed a publicly available RNA-Seq dataset of A549 lung epithelial cells infected with SARS-CoV-2 (Blanco-Melo et al. 2020) to investigate the effect of viral infection on gene and RNA isoform expression (Fig. 1A). The dataset includes wild-type (WT) or ACE2 overexpressing (ACE2) A549 cells infected with SARS-CoV-2 at a low (0.2) or high (2.0) multiplicity of infection (MOI) for 24 hours (Fig. 1A).

**Figure 01.**
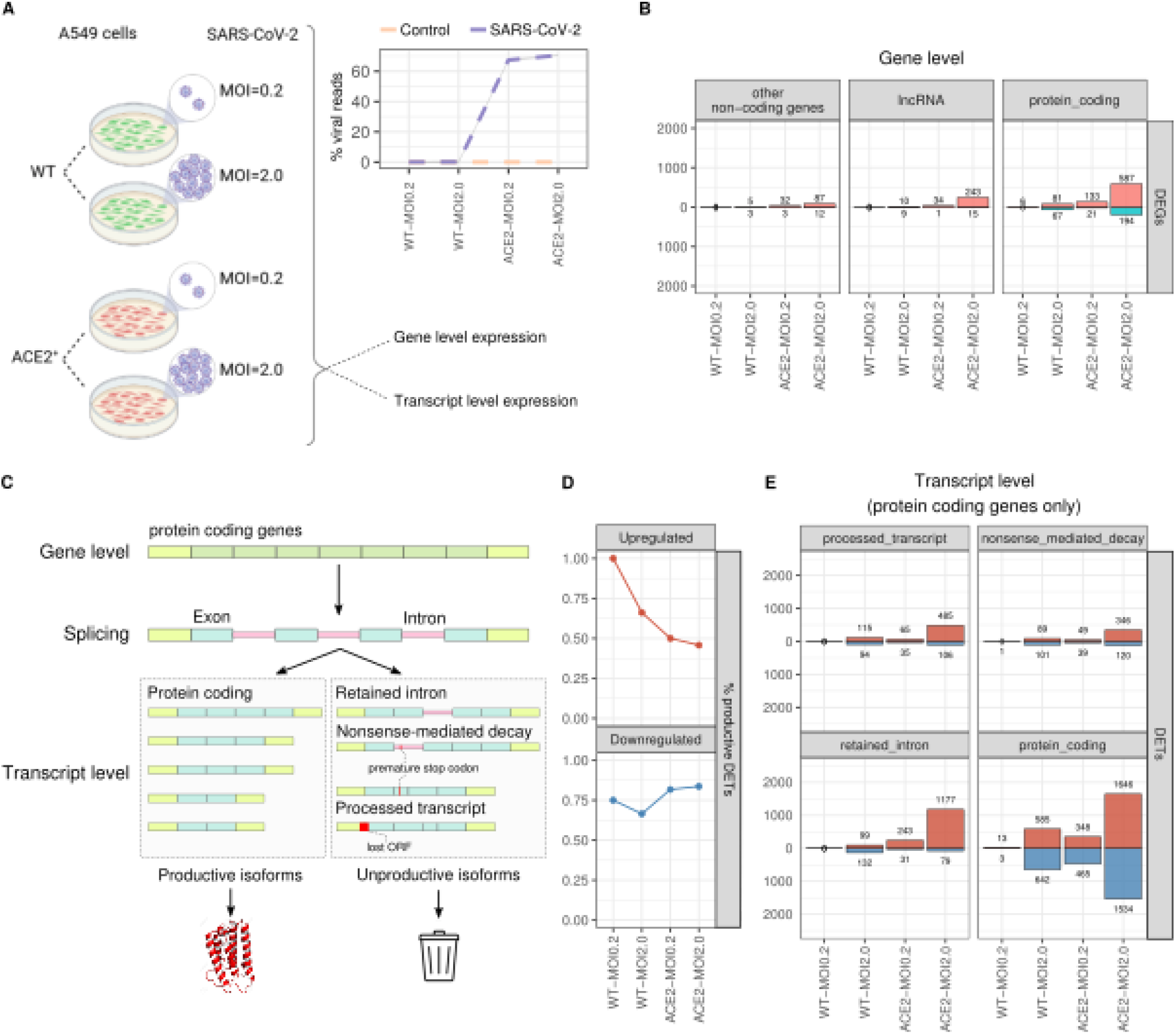
SARS-CoV-2 promotes gene and transcript-level perturbations according to the viral load. **A.** Experimental design of the RNA-Seq dataset obtained from (Blanco-Melo et al. 2020). A549 WT or ACE2 overexpressing cells were submitted to SARS-CoV-2 infection at a MOI of 0.2 or 2.0 for 24h. **B.** Number of up- and downregulated DEGs in each gene category: other non-coding genes, lncRNA genes and protein-coding genes. DEGs were considered those with an adjusted p-value < 0.05 and abs(log_2_FoldChange) > 2. **C.** Human genes can be spliced into two main transcript categories: productive (protein-coding) and unproductive (retained intron, NMD, and processed transcripts) splicing isoforms. Unproductive isoforms are not canonically translated into proteins. **D.** Proportion of productive isoforms among the up- and downregulated DETs of protein-coding genes in each cell. **E.** Number of up- and downregulated DETs of each transcript type of protein-coding genes (intron retention, NMD, processed transcript, and protein-coding) in each cell. DETs were considered those with an adjusted p-value < 0.05 and abs(log_2_FoldChange) > 2.

Consistent with viral load results (Fig. 1A), which was higher in ACE2 cells, we detected the highest number of differentially expressed genes (DEGs) in ACE2-MOI2.0 cells (Fig. 1B and Table S1). In these cells, we identified 243 up- and 15 downregulated long non-coding RNAs (lncRNAs), 87 up- and twelve downregulated non-coding RNAs (ncRNAs), and 587 upregulated and 194 downregulated protein-coding DEGs (Fig. 1B). The total number of DEGs in ACE2-MOI2.0 cells was markedly higher than those in ACE2-MOI0.2, WT-MOI2.0, and WT-MOI0.2 cells (Fig. 1B).

Gene-level expression analysis is standard for RNA-Seq but often overlooks alternative splicing into productive or unproductive isoforms (Fig. 1C), a mechanism cells use for dynamic protein expression control (Sibley 2014). For example, the abundance of productive isoforms of inflammatory genes in response to tumor necrosis factor (TNF) is mostly regulated through splicing, not transcription (Hao & Baltimore 2013).

We demonstrated that the proportion of differentially expressed productive transcripts (DETs) of protein-coding genes decreases among upregulated DETs in ACE2 overexpressing cells and is also affected by higher MOI (Fig. 1D). Interestingly, the number of upregulated processed transcript and NMD DETs was higher in WT-MOI2.0 cells than that seen in ACE2-MOI0.2 cells (Fig. 1E), which was not detected at the gene-level (Fig. 1B) or for retained intron isoforms (Fig. 1E). Transcript-level analysis reveals a more detailed landscape of how genotype (WT or ACE2) and MOI (high or low) might distinctly influence the expression of transcripts in some isoform categories, but not others, in SARS-CoV-2 infected cells.

As ACE2-MOI2.0 cells presented the highest number of DEGs and DETs, we focused our subsequent analyses on those cells. Results for other cells are available in the supplementary material (Fig. S1).

### ARS-CoV-2 selectively induces unproductive isoforms of IFN signaling genes while promoting productive expression of inflammatory genes

We performed a gene set enrichment analysis (GSEA) on differentially expressed genes (DEGs) and differentially expressed transcripts (DETs) in infected cells to elucidate the biological functions modulated by SARS-CoV-2. Utilizing both productive and unproductive transcripts of protein-coding genes in ACE2-MOI2.0 cells, we examined the impact of each component on selected pathways (Fig. 2A). Pathways implicated in pathogen response were chosen based on manual curation of the enrichment analysis and are illustrated in Figure 2. Comprehensive GSEA results can be found in the supplementary material (Table S2).

**Figure 02.**
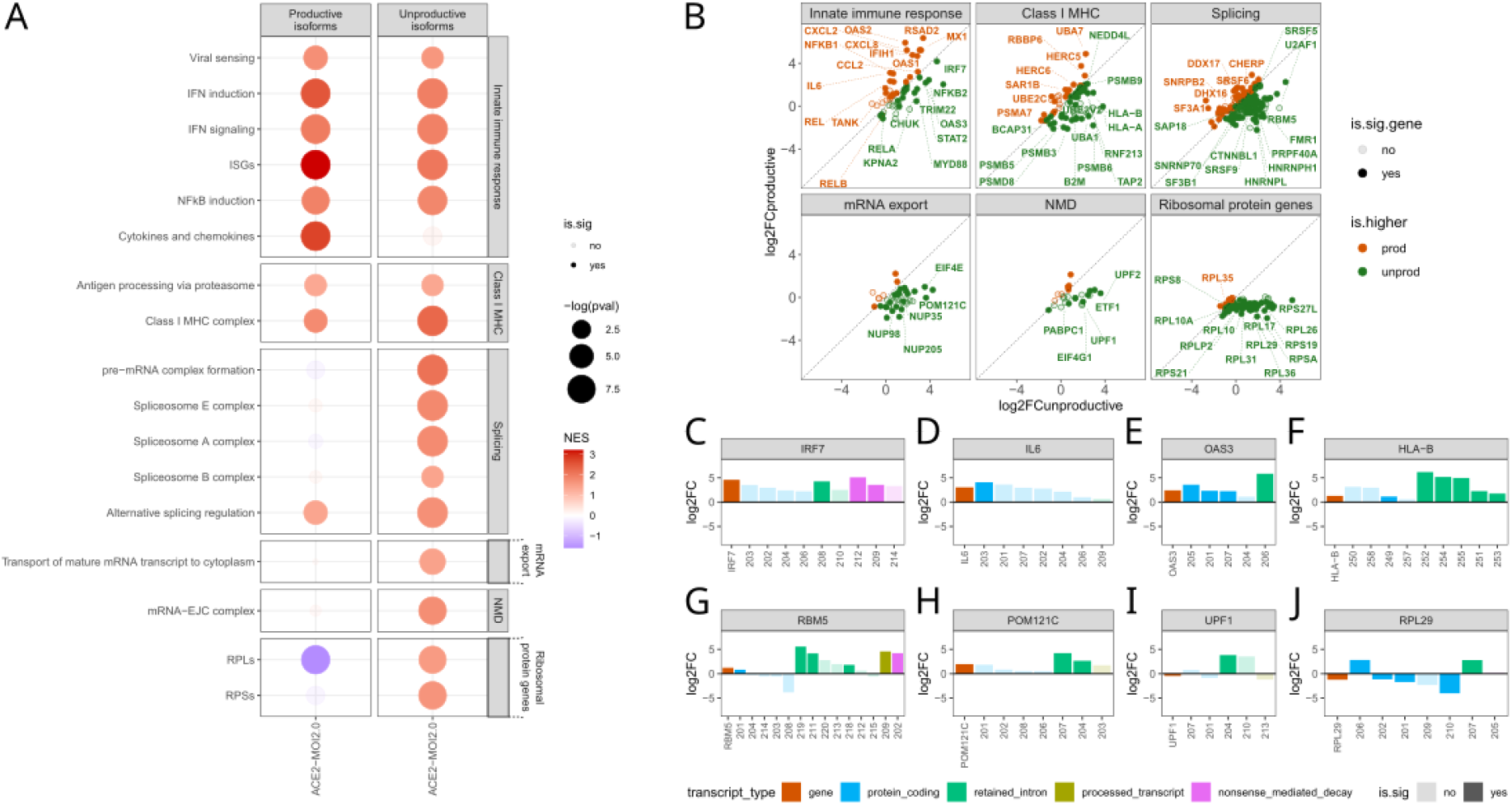
SARS-CoV-2 induces unproductive splicing isoforms of IFN signaling, class I MHC, and splicing genes. **A.** GSEA results for the productive and unproductive DETs in ACE2-MOI2.0 cells. The size of the dots is proportional to the -log_10_pvalue of the *fgsea* enrichment for each custom pathway and the color is related to the normalized enrichment score (NES - red positive, blue negative). **B.** Aggregated log2FoldChange of productive versus unproductive isoforms of the genes in the pathways depicted in **A**. For each gene, the aggregate log2FoldChange of each type of isoform was calculated as the mean shrunk log2FoldChange value of all productive or unproductive isoforms of the gene. The diagonal dotted line separates genes which had a higher aggregate productive fold change (red dots) from those that had a higher aggregate unproductive fold change (blue dots). Gene not significantly DE at the gene-level are colored in a transparent hue. **C-J.** Gene (red) and transcripts (blue=protein-coding, green=retained intron, gold=processed transcript, and purple=NMD) log2FoldChange and significance status of *IRF7* (C), *IL6* (D), *OAS3* (E), *HLA-B* (F), *RBM5* (G), *POM121C* (H), *UPF1* (I), and *RPL29* (H) in ACE2-MOI2.0 cells.

We observed an upregulation in IFN response-related pathways, such as viral sensing, IFN induction, IFN signaling, and interferon-stimulated genes (ISGs) in both productive and unproductive isoforms (Fig. 2A) and at the gene level (Fig. S1A). However, key genes in the IFN-mediated antiviral response, including MYD88, STAT2, and IRF7, demonstrated a higher fold change in unproductive isoforms (Fig. 2B). Notably, the IRF7 gene expressed only unproductive overexpressed isoforms in ACE2-MOI2.0 cells (Fig. 2C and Fig. S1C), whereas WT-MOI0.2 cells predominantly overexpressed productive IRF7 transcripts (Fig. S1D). We propose that SARS-CoV-2 may amplify a constitutive feedback mechanism involving intron 4 retention in the Irf7 gene seen in mice, which suppresses productive IRF7 isoforms expression and dampens the IFN response following innate immune stimulation (Frankiw et al. 2019).

ISGs exhibited a more balanced overexpression of productive and unproductive isoforms compared to early IFN signaling genes (Fig. 2B). Key genes, such as MX1, OAS1, OAS2, and RSAD2, displayed a higher fold change in productive isoforms (Fig. 2B), whereas TRIM22 and OAS3 (Fig. 2D) showed higher expression of unproductive isoforms. A recent study demonstrated that two common human OAS1 locus variants (rs1131454 and rs10774671) are enriched in European and African COVID-19 patients requiring hospitalization (Banday et al. 2022). These variants elevate the expression of an OAS1 transcript with a premature stop-codon, resulting in transcript NMD degradation, reduced OAS1 protein expression, and impaired viral clearance (Banday et al. 2022). In a similar vein, Frankiw et al., 2020, reported that splicing at a frequently used splice site in the mouse Oas1g gene generates an NMD-targeted transcript, while removal of the site leads to increased Oas1g expression and an enhanced antiviral response (Frankiw et al. 2020). Hao and Baltimore, 2013, showed that TNF stimulation-induced innate immune gene expression is regulated by splicing, particularly in the regulation of productive isoform expression, rather than transcription (Hao & Baltimore 2013).

These findings support our results, emphasizing the significance of the productive versus unproductive balance in regulating key ISGs in response to SARS-CoV-2. Interestingly, gene- and transcript-level ISG expression was increased only in cells with the highest (ACE2-MOI2.0) and lowest (WT-MOI0.2) viral loads (Fig. S1A). This suggests that SARS-CoV-2 can silence ISG expression in cells with intermediate viral loads but not in cells infected with either an excess or a paucity of viral particles. Blanco-Melo et al., 2020, proposed that this effect, which they also observed for IFNɑ and IFNβ proteins, could stem from a stoichiometric competition between host and viral components, favoring the virus in cells with intermediate viral loads. When few viral particles are present, host components outnumber viral ones, leading to a successful response. Conversely, the presence of numerous viral copies may induce a robust host response, potentially shifting the balance in favor of the host (Blanco-Melo et al. 2020). Our findings lend support to this notion for ISG expression in infected cells as well.

Taking a different perspective, numerous genes involved in promoting and regulating inflammation via NFκB, such as TANK, REL, RELB, and NFKB1, displayed a higher fold change in their productive isoforms (Fig. 2B). Conversely, the gene CHUK, which encodes for the NFκB Inhibitor Kinase Alpha (IKK-ɑ), exhibited a greater overexpression of unproductive isoforms (Fig. 2B), potentially contributing to increased NFκB-mediated inflammation. Additionally, inflammatory cytokine and chemokine genes, including IL6, CXCL8, CXCL2, and CCL2 (Fig. 2B and 2E), demonstrated a marked positive enrichment of productive isoforms (Fig. 2A). No enrichment was detected for the unproductive isoforms of cytokine and chemokine genes in ACE2-MOI2.0 cells (Fig. 2A).

These findings align with a clinical hallmark of severe COVID-19: exacerbated lung inflammation driven by hyperresponsive immature neutrophils and classical monocytes (COvid-19 Multi-omics Blood ATlas (COMBAT) Consortium 2022). Our results suggest that the critical imbalance observed in COVID-19 patients may result from the biased impact of SARS-CoV-2 on inducing unproductive splicing of IFN-related genes, an effect not seen for key inflammatory genes.

### SARS-CoV-2 Promotes Unproductive Splicing of Class I MHC Genes

Pathways associated with antigen presentation through class I MHC exhibited greater upregulation among unproductive isoforms compared to productive isoforms (Fig. 2A). Crucial antigen presentation genes with higher unproductive fold changes included proteasome components PSMB3, PSMB5, PSMB6, PSMB9, and PSMD8. Genes encoding core class I MHC complex elements, such as B2M, HLA-A, and HLA-B, also displayed a higher unproductive fold change. The HLA-B gene revealed five distinct overexpressed retained intron isoforms in ACE2-MOI2.0 cells (Fig. 2F and Fig. S1C). This upregulation of unproductive isoforms could potentially decrease HLA-B protein production in SARS-CoV-2 infected cells, impairing antigen presentation.

A 10bp deletion in the 3’ region of intron 1 of the HLA-B*1501 allele was identified in a bone marrow donor, promoting intron retention and resulting in the absence of HLA-B15 antigen from the donor’s serum (Curran et al. 1999). Similarly, melanoma cells possess a mutation at the 5’ splice donor site of intron 2 of the HLA-A2 gene, causing the absence of the HLA-A2 antigen and likely protecting these cancerous cells from immune surveillance (Wang et al. 1999). Our findings indicate that SARS-CoV-2 infection may also lead to the loss of class I MHC antigens due to increased intron retention, potentially rendering infected cells incapable of presenting viral antigens to the immune system. Reduced expression of HLA-DR in monocytes represents a molecular hallmark of severe COVID-19 (COvid-19 Multi-omics Blood ATlas (COMBAT) Consortium 2022). Other studies have demonstrated that SARS-CoV-2 ORF6 inhibits class I MHC production by blocking the STAT1-IRF1-NLCR5 axis (Yoo et al. 2021) and that ORF3a and ORF7a downregulate class I MHC surface expression in infected cells through distinct mechanisms (Arshad et al. 2022). Our results expand on this existing evidence by revealing that SARS-CoV-2 also promotes unproductive splicing of key class I MHC genes.

### Unproductive Isoforms of Splicing Factors and Mediators Highly Expressed During SARS-CoV-2 Infection

In ACE2-MOI2.0 cells, significant upregulation of several unproductive transcripts of genes related to various stages of the splicing mechanism was observed (Fig. 2A). Key genes encoding proteins involved in recognizing splicing elements for splice site selection, such as HNRNPH1 and HNRNPHL (Fig. 2B), and those regulating alternative splicing, including RBM5 (Fig. 2G) and SRSF5, displayed a higher fold change of their unproductive isoforms in ACE2-MOI2.0 cells (Fig. 2B). Cells often modulate splicing factor abundance by regulating unproductive isoform expression, subsequently affecting the splicing of several other genes in response to stress (Ding et al. 2020).

SARS-CoV-2 and numerous other viruses exploit this mechanism by hijacking, disrupting, or modulating host splicing machinery. For instance, the herpes simplex virus type 1 protein ICP27 binds to SRPK1 and SR proteins, inhibiting cellular splicing (Sandri-Goldin 2008), while the Epstein-Barr virus SM protein hijacks the cellular protein SRp20 during alternative splicing (Verma et al. 2010). The yellow fever virus NS5 protein interacts with proteins involved in polyadenylation (Bronzoni et al. 2011), and the influenza A NS1 protein interacts with the U6 snRNA, inhibiting pre-mRNA splicing (Qiu et al. 1995). Our findings reveal that SARS-CoV-2 selectively manipulates the host’s splicing machinery to induce unproductive transcript expression for a subset of essential antiviral genes and genes associated with the splicing machinery itself. Wang et al. (2022) recently demonstrated that COVID-19 patients exhibit global alterations in alternative splicing, and disease severity correlates with the degree of splicing perturbation (Wang et al. 2022).

### Nuclear mRNA Export and NMD Pathways Impacted by SARS-CoV-2 Induction of Unproductive Isoform Expression

Genes involved in transporting mature mRNA transcripts to the cytoplasm, such as NUP35, NUP98, NUP205, and POM121C (Fig. 2B and Fig. 2H), exhibited higher upregulation of their unproductive isoforms (Fig. 2A and 2B). SARS-CoV-2 ORF6 protein has been consistently shown to interact with and block the nuclear pore, hindering the entry of IFN mediators into the nucleus and the exit of mature ISG mRNAs from the nuclear compartment (Addetia et al. 2021; Gordon et al. 2020; Kato et al. 2021; Miorin et al. 2020). Our findings indicate that core nuclear pore genes are also affected at the transcript level through the induced expression of unproductive isoforms by SARS-CoV-2.

Similarly, numerous genes involved in the NMD pathway exhibited a shift toward an unproductive expression profile, including genes of the mRNA-EJC complex formation, the starting point of the NMD pathway (Fig. 2A and 2B). Although the UPF1 gene was downregulated at the gene level, it displayed increased expression of a retained-intron isoform at the transcript level (Fig. 2I). The protein encoded by UPF1 is essential for recognizing truncated transcripts and initiating the NMD process. NMD-involved proteins can also detect and degrade viral RNA, serving as an additional layer of antiviral response (Leon & Ott 2021; Sarkar et al. 2022). Key genes in NMD, such as ribosomal protein (RP) genes, were also distinctly affected in their productive and unproductive isoforms by SARS-CoV-2 infection (Fig. 2A and 2B). Genes coding for RPs from both large and small subunits, including RPS19, RPL26, and RPL29 (Fig. 2J), demonstrated reduced expression of multiple productive isoforms and overexpression of unproductive transcripts (Fig. 2B).

Given that SARS-CoV-2 NSP16 protein directly disrupts splicing (Banerjee et al. 2020), we anticipated detecting compromised splicing in various genes. Our results confirmed this hypothesis but revealed that the effect is not evenly distributed across different genes. In addition to splicing and NMD genes, unproductive splicing preferentially occurs in genes of the IFN-mediated immune response and class I MHC complex, while it is virtually absent in cytokine and chemokine genes.

### Genes with upregulated unproductive transcripts are less translated in SARS-CoV-2 infected cells

Given the substantial impact of SARS-CoV-2 infection on the expression of unproductive splicing isoforms of key genes, we investigated whether these alterations could also be detected at the protein level. We analyzed a publicly available translatome dataset of colorectal adenocarcinoma cells (Caco-2) infected with SARS-CoV-2 at an MOI of 1.0 at 2-, 6-, 10-, or 24-hours post-infection (Fig. 3A; Bojkova et al. 2020). Nascent peptides were labeled at each time point using heavy isotope-containing lysine (K8) and arginine (R10), enabling their quantification by LC-MS/MS (Bojkova et al. 2020) (Fig. 3A). The number of differentially translated proteins (DTPs) in response to SARS-CoV-2 infection steadily increased from 2 to 24 hours (Fig. 3B), with the majority being downregulated, except at 2h post-infection (Fig. 3B). Among the 606 DTPs found at any time point (Table S3), 237 also had differentially expressed transcripts (DETs) in ACE2-MOI2.0 cells (Fig. 3C). These genes with both DETs and DTPs exhibited a larger fold change of their unproductive isoforms (Fig. 3C) and were highly represented among downregulated proteins at 24h in the translatome (Fig. 3C). A potential caveat is that we also observed a higher number of underrepresented proteins compared to overrepresented ones among the productive isoforms of DETs, which suggests that the downregulation may not be solely attributed to the unproductive isoforms (Fig. 3C).

**Figure 03.**
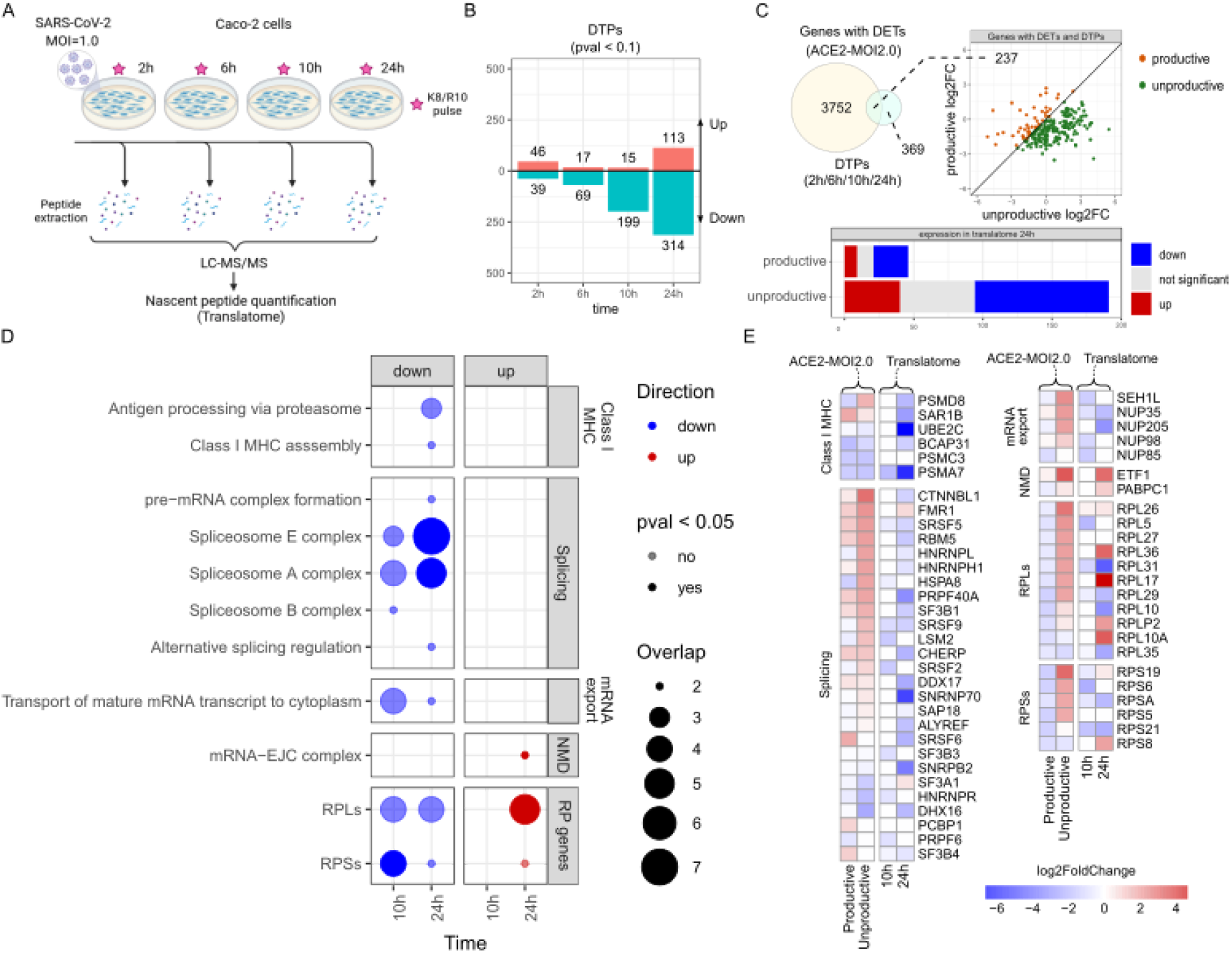
Genes with upregulated unproductive transcripts also have lower translation rate in SARS-CoV-2 infected cells. **A.** Experimental design of the translatome dataset obtained from (Bojkova et al. 2020). Caco-2 cells were infected with SARS-Cov-2 at an MOI of 1.0. Nascent peptides were labeled with isotope-containing lysine (K8) and arginine (R10), which allowed for their quantification by LC–MS/MS. Differential translation rate was calculated by comparing infected versus mock infected cells in each time point. **B.** Number of up- (red) and downregulated (blue) DTPs in each time point. DTPs were those with a p-value < 0.1. **C.** Number of genes that have DTPs in any time point in the translatome and also DETs in the RNA-Seq of ACE2-MOI2.0 cells. Genes with DETs and DTPs according to the log2FoldChange of their productive and unproductive isoforms. Number of genes with DETs and DTPs according to their differential expression in the translatome at 24h. **D.** Overrepresentation analysis of up- (red) and downregulated (blue) DTPs using custom gene sets. Significant pathways were those with p-value < 0.05. The size of the points is proportional to the number of DTPs that are also in each pathway. **E.** Heatmap depicting the aggregate log2FoldChange of the productive and unproductive transcripts (left columns) of genes that are also DTPs at 10h or 24h in the translatome (right columns).

We performed an overrepresentation analysis (ORA) with the up- and downregulated DTPs in each time point to verify which pathways were the most impacted by SARS-CoV-2 at the translatome level (Fig. 3D). Enrichment results were more significant at 10h and 24h. Proteins related to antigen processing via the proteasome and assembly of the class I MHC complex were downregulated at 24h (Fig. 3D), including PSMD8 and SAR1B (Fig. 3E). The genes that code for those proteins also presented a positive fold change of their unproductive isoforms in ACE2-MOI2.0 cells in the RNA-Seq (Fig. 3E). HLA-A and B2M, core class I MHC components which also had increased expression of unproductive isoforms at the transcriptome level, had a negative fold change at the translatome level, but with a non-significant associated adjusted p-value (Table S3). The same occurred with the proteins STAT1 and RELA (Table S3), which are key transducers of the IFN-mediated antiviral response. Spliceosome E and A complex proteins were also less translated at 10h and 24h (Fig. 3D). In total, 16 out of 25 splicing proteins that had a reduced translation rate at 10h or 24h also presented a higher fold change of unproductive isoforms of their corresponding gene in the RNA-Seq, including RBM5, HNRPNH1, and HNRNPL (Fig. 3E). Nuclear pore complex proteins (NUPs) involved in the transport of mature mRNA transcripts to the cytoplasm were also less translated at 10h and 24h in Caco-2 cells infected with SARS-CoV-2 (Fig. 3D). Among those, NUP35, NUP205, and NUP98 presented a positive fold change of unproductive isoforms in the transcriptome (Fig. 3E). Contrary to what we had expected, proteins of the mRNA-EJC complex of the NMD pathway were upregulated in the translatome (Fig. 3D), including ETF1 and PABPC1, which also showed overexpression of their unproductive isoforms in the RNA-seq (Fig. 3E). Ribosomal proteins involved in the NMD process, on the other hand, were mostly downregulated in the translatome (Fig. 3D), including RPL29, which also had downregulated productive isoforms and upregulared unproductive transcripts in the RNA-seq (Fig. 3E).

The authors who showed that the SARS-CoV-2 NSP16 protein disrupts splicing also demonstrated that the NSP1 viral protein binds to host ribosomes and inhibits translation (Banerjee et al. 2020). Our results suggest that the effects of these viral proteins are not generalist, but rather lead to splicing disruption and translational silencing of transcripts involved in specific biological functions, namely the IFN-mediated innate immune response, class I MHC antigen presentation, splicing, and mRNA processing.

### SARS-CoV-2 proteins interact with key antiviral, splicing, and nuclear transport proteins

Lastly, we analyzed networks of experimentally defined physical interactions between SARS-CoV-2 proteins and host proteins (Gordon et al. 2020; Stukalov et al. 2021). The first protein-protein interaction network (PPI) had 23 viral proteins with significant interactions with 330 host target proteins (Gordon et al. 2020). The second PPI included 22 SARS-CoV-2 proteins targeting 876 host proteins (Stukalov et al. 2021).

Several SARS-CoV-2 proteins interfere with the host’s innate immune system (Fig. 4A). ORF7b interacts with IL10RB and the MAVS protein (Fig. 4B), a key transducer of the signal from viral sensing elements such as RIG-I and MDA5 (Chen et al. 2021). NSP13 binds to the TBK1 protein (Fig. 4B), another transducer of the viral sensing signaling pathway (Zhang et al. 2022), and ORF3 interacts with IL10RB and IFNGR1 (Fig. 4B), the main IFN-gamma receptor. The M protein interacts with both the IRAK1 protein (Fig. 4B), a transducer of toll-like receptor-mediated responses (Bruni et al. 2015), and JAK2 (Fig. 4B), a mediator of the IFN receptor signaling. None of the ISGs that were altered at the isoform and translation levels appeared as direct targets of SARS-CoV-2 proteins (Fig. 4B), suggesting that the virus deploys distinct, complementary mechanisms to interfere with the innate antiviral host response mediated by interferons. RIPK1 was the only host protein related to the NFκB-mediated inflammatory signaling targeted by a viral protein, NSP12 (Fig. 4D). Moreover, we did not find any interaction between viral proteins and cytokine or chemokine proteins (Fig. 4B-D).

**Figure 04.**
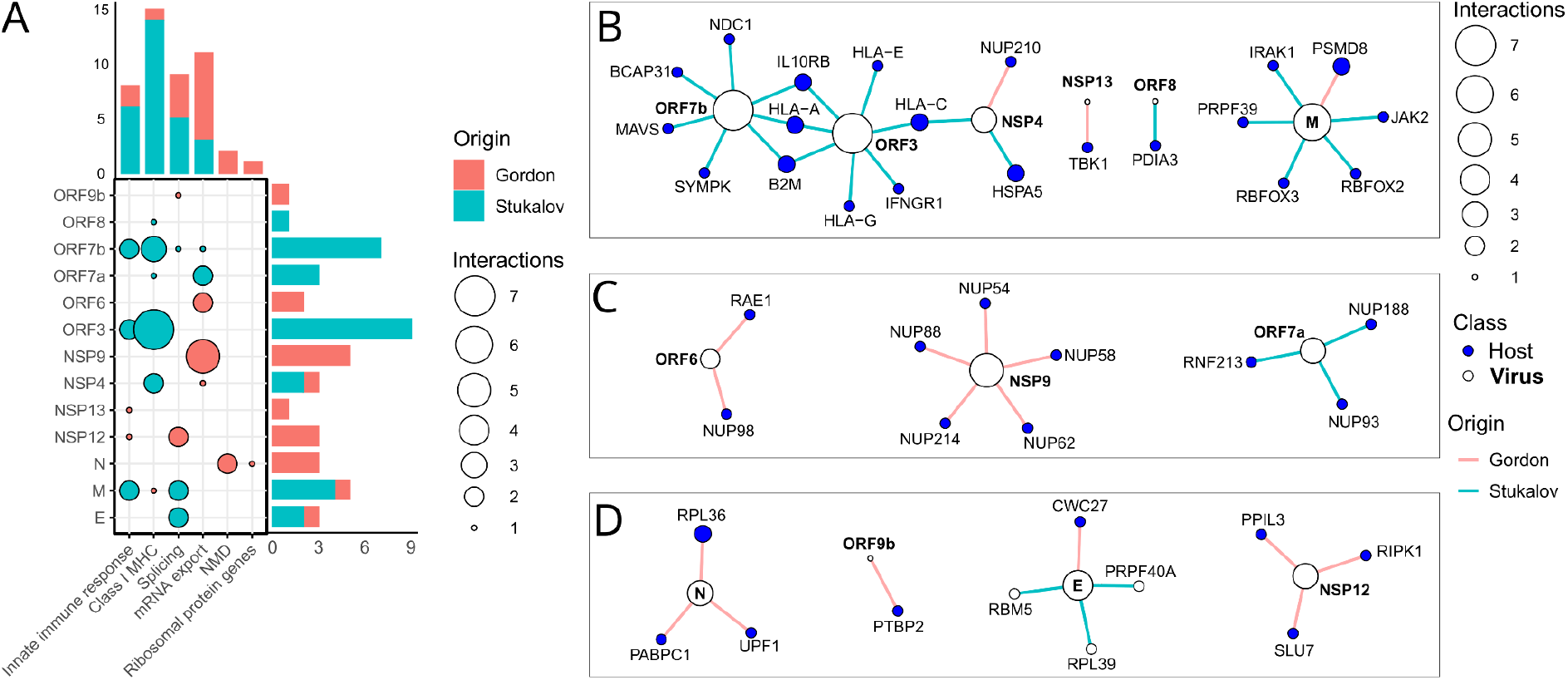
SARS-CoV-2 proteins interact with key host antiviral, splicing and nuclear export RNAs and proteins. **A.** Number of host proteins (Gordon and Stukalov PPIs) in each pathway that interacts with SARS-CoV-2 proteins. **B.** Virus versus host interaction network depicting SARS-CoV-2 proteins (white nodes) and proteins (blue nodes) they interact with. The color of the edges depicts the dataset in which the association was detected: pink = Gordon et al., lightblue = Stukalov et al. The size of the nodes is proportional to the number of connections they make.

SARS-CoV-2 peptides also heavily target host antigen presentation and class I MHC proteins (Fig. 4A). The NSP4 viral protein targets the HLA-C and HSPA5 proteins and ORF3 targets HLA-A, HLA-C, HLA-E, HLA-G, and the B2M protein (Fig. 4B). ORF7b also interacts with the HLA-A and B2M proteins (Fig. 4B). HLA-A and B2M are core components of the class I MHC complex (Fisette et al. 2020) which also had an increased expression of several retained intron transcripts in the RNA-Seq (Fig. 2B). ORF8 binds to PDIA3 (Fig. 4B), a protein involved in antigen loading to the class I MHC complex (Fisette et al. 2020) which also presented strong upregulation of one of its unproductive isoforms, PDIA3-204, in ACE2-MOI2.0 cells (Table S1). The viral protein M interacts with the proteasome protein PSMD8 (Fig. 4B). The *PSMD8* gene also showed strong overexpression of unproductive splicing isoforms and reduced translation in SARS-CoV-2 infected cells (Fig. 2B and Fig. 3E). Thus, analysis of the virus-host interactome shows that SARS-CoV-2 proteins interfere with the presentation of viral antigens at distinct levels, through both direct (protein-protein interactions) and indirect (alternative splicing) manipulation of key proteins of the antigen presentation machinery.

Export of mature mRNA transcripts to the cytoplasm was the process with the most host peptides targeted by SARS-CoV-2 proteins (Fig. 4A). Several NUPs and mRNA export proteins were targeted by the viral proteins NSP4, NSP9, ORF6, ORF7a, and ORF7b, including RAE1, and NUP98 (Fig. 4C). In the early days of the COVID-19 pandemic, (Gordon et al. 2020), argued that the interaction between ORF6 and the host proteins NUP98 and RAE1 might prevent the export of IFN-stimulated transcripts from the nucleus. Later studies confirmed that hypothesis, showing that ORF6 functions as a potent inhibitor of the IFN-signaling pathway by preventing several key antiviral transducers, such as STAT1, IRF3 from entering the nucleus, and ISG transcripts from leaving the nuclear envelope (Addetia et al. 2021; Frieman et al. 2007; Kato et al. 2021; Miorin et al. 2020). Proteins of several other viruses interact with host nuclear pore proteins during the viral replication cycle, including the HIV-1 capsid protein (Di Nunzio et al. 2013; Matreyek et al. 2013) and the NS3 protein of the Zika and Dengue viruses (De Jesús-González et al. 2020).In conclusion, we showed that SARS-CoV-2 affects NUPs and other mRNA nuclear export genes also at the transcriptional and translational levels, increasing the levels of unproductive splicing isoforms of these genes and, therefore, reducing protein translation.

Core splicing machinery proteins are also targeted by SARS-CoV-2 proteins (Fig. 4A). The E protein interacts with PRPF40A, CWC27, and RBM5 (Fig. 4D), all of which participate in the early steps of RNA splicing. The *RBM5* gene also presented increased expression of unproductive transcripts in SARS-Cov-2 infected cells (Fig. 2G). Similarly, the hepatitis delta virus (HDV) promotes unproductive *RBM5* expression due to the disrupting interaction of the HDV genomic RNA with the splicing factor SF3B155 (Tavanez et al. 2020). The SARS-CoV-2 M protein interacts with PRPF39, a component of the U1snRNP complex, and with the alternative splicing regulators RBFOX2 and RBFOX3 (Fig. 4D). RBFOX2 is a hub protein in a network connecting splicing factors to differentially spliced genes, including *STAT1* and *NUP43*, in herpes virus-induced liver hepatocarcinoma patients (Cai et al. 2020). In mice, an *Rbfox3* isoform promotes the inclusion of an alternative exon containing a premature stop-codon in *Rbfox2* transcripts, leading to their degradation via NMD (Dredge & Jensen 2011). Three other SARS-CoV-2 proteins, NSP12, ORF7b, and ORF9b physically interact with the splicing factors PPIL3 and SLU7, SYMPK, and PTBP2, respectively. These results suggest that the splicing perturbations promoted by SARS-CoV-2 characterized here might be a result of a combination of multiple viral factors acting together to disrupt host RNA processing, rather than the sole effect of the NSP16 sequestering U2 snRNA described by Banerjee et al. 2020.

Two SARS-CoV-2 proteins, N and E, interact with key NMD proteins. The N protein interacts with both UPF1 and PABPC1 (Fig. 4D). The rotavirus protein NSP5 also targets UPF1, leading to its proteasomal degradation and facilitating viral replication due to a burdening of the NMD pathway (Sarkar et al. 2022). The porcine epidemic diarrhea virus (PEDV) reduces *PABPC1* expression, while overexpression of the gene refrains viral replication (Wu et al. 2022). Interestingly, similarly to SARS-CoV-2, the PEDV nucleocapsid (N) protein also binds to PABPC1 (Wu et al. 2022). The NMD pathway works as a complementary mechanism of antiviral response by promoting viral RNA decay, at the same time as host unproductive RNAs are also degraded (Mailliot et al. 2022). Our results demonstrate that SARS-CoV-2 physically targets core proteins of this mechanism, while also promoting unproductive expression of the genes that code for those proteins.

## Conclusions

The central finding of our study is that SARS-CoV-2 disrupts the host’s splicing of several genes to potentially silence the antiviral responses of infected cells (Fig. 05). After entering a human cell, SARS-CoV-2 expresses NSP16, which translocates to the nucleus, where it binds to and sequesters snRNAs U1 and U2 (Fig. 5A; Banerjee et al. 2020). This leads to a disruption in splicing, which increases the expression of intron containing transcripts, as well as other unproductive transcripts, such as those containing premature stop codons and those missing an ORF (Fig. 5A). These transcripts, contrary to their productive counterparts, do not become translated into canonical proteins in the cytoplasm (Fig. 5A). Here we showed that this phenomenon heavily affects genes involved in the innate immune response against viruses, including the interferon-mediated response. Surprisingly, cytokine and chemokine genes are not consistently affected, overexpressing mostly productive splicing isoforms, which can be translated into proteins (Fig. 5B). Viral antigens are mostly presented through the class I MHC pathway, which is also affected by the disruption of splicing promoted by SARS-CoV-2 (Fig. 5C). Key genes involved in this pathway are less translated in infected cells and are targeted by several viral proteins. This likely helps the virus to avoid detection by the adaptive immune system. Lastly, SARS-CoV-2 also induces the expression of unproductive splicing isoforms of genes from the splicing machinery, as well as genes related to the NMD pathway and to the export of mature mRNAs from the nucleus (Fig. 5D-F). Several genes in these pathways also present reduced translation in infected cells and produce proteins or RNAs that are directly targeted by viral proteins. This shows that SARS-CoV-2 disrupts splicing and mRNA processing using multiple resources both at the start of the process as well as downstream.

**Figure 05.**
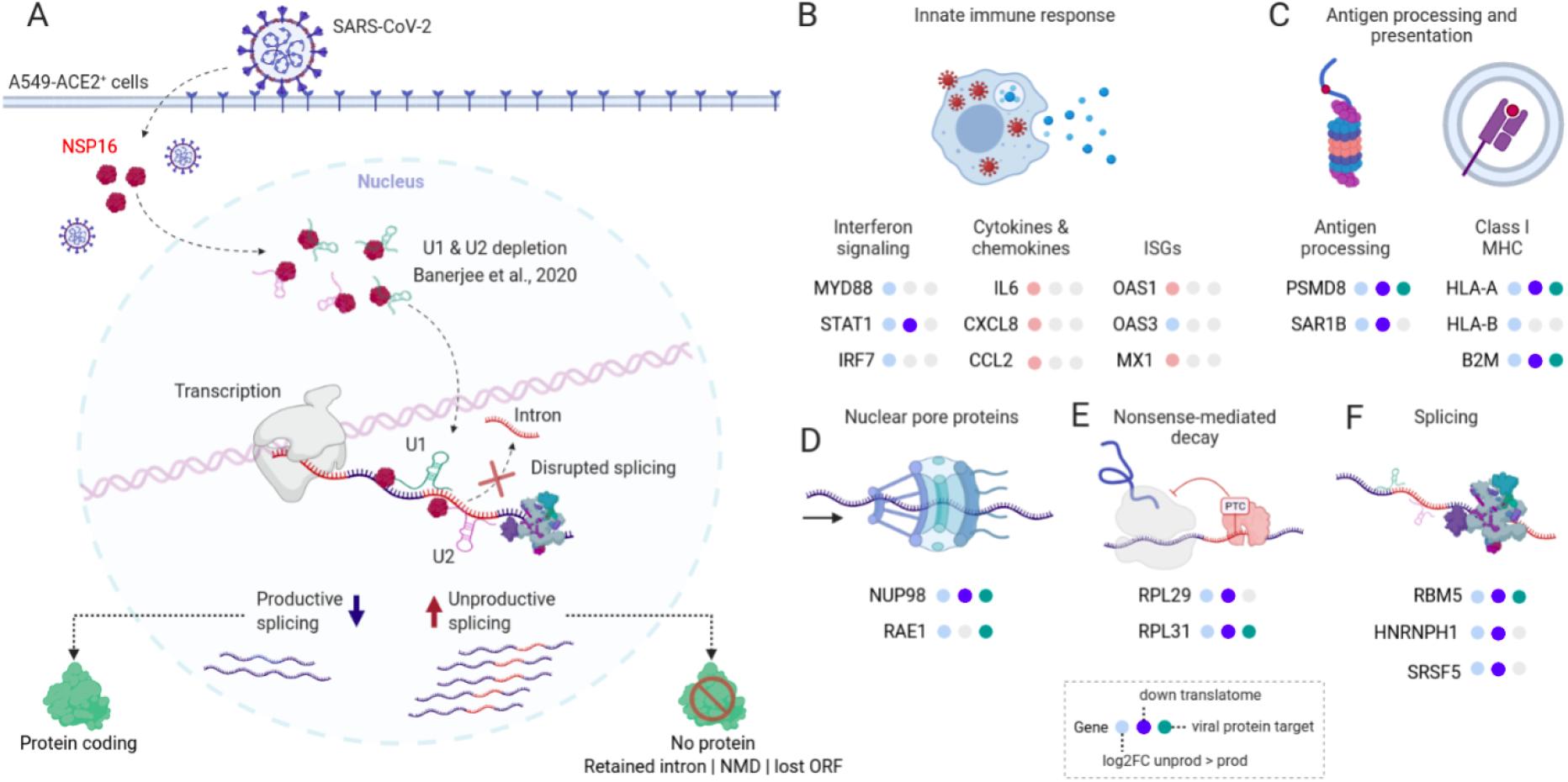
SARS-CoV-2 disrupts splicing in infected cells and induces the expression of unproductive isoforms of interferon, class I MHC and splicing genes. **A.** SARS-CoV-2 NSP16 protein binds to U1 and U2 snRNAs and disrupts splicing (Banerjee et al. 2020). This leads to increased expression of unproductive splicing isoforms (retained intron, NMD, lost ORF), which are not translated into proteins. **B-F.** Genes involved with the innate immune response (**B**), class I MHC antigen processing and presentation (**C**), nuclear pore proteins (**D**), NMD (**E**), and splicing (**F**) are affected by SARS-CoV-2 at the transcriptome, translatome and interactome levels. Color code: light blue - genes with a higher log_2_ fold change of unproductive isoforms, pink - genes with a higher log_2_ fold change of productive isoforms, blue - proteins with a reduced translation rate, green - proteins or RNAs targeted by a viral protein, gray - no alterations in translatome or interactome layers.

## Methods

### Public datasets

Raw RNA-Seq reads were obtained from the study of (Blanco-Melo et al. 2020) (GSE147507). These reads correspond to triplicate samples of WT A549 cells infected with SARS-CoV-2 at an MOI of 0.2 (series 2) or 2.0 (series 5), ACE2 overexpressing cells infected with SARS-CoV-2 at an MOI of 0.2 (series 6) or 2.0 (series 16), and triplicate samples of the corresponding mock-infected cells in each series. Cells were infected for 24 hours before RNA extraction. Libraries were prepared with the TruSeq RNA Library Prep Kit v2 (Illumina) or TruSeq Stranded mRNA Library Prep Kit (Illumina) and sequenced on the Illumina NextSeq 500 platform (Blanco-Melo et al. 2020).

The differential translation rate results table was obtained from the supplementary material of (Bojkova et al. 2020). These translatome results were obtained in Caco-2 cells infected with SARS-CoV-2 (MOI=1.0) for 2, 6, 10, or 24 hours and compared to equivalent mock-infected cells. Nascent proteins were labeled at each time point using heavy isotope-containing lysine (K8) and arginine (R10), which allowed for their quantification by LC–MS/MS. Differentially translated proteins in each time point were those with p-value < 0.1.

Protein-protein interaction networks were obtained from the supplementary materials of (Gordon et al. 2020) and (Stukalov et al. 2021), and the protein-RNA interaction network was obtained from the supplementary material of (Banerjee et al. 2020). The protein-protein interaction networks were obtained from HEK-293T (Gordon et al. 2020) or A549 (Stukalov et al. 2021) cells individually transfected with expression vectors each containing one SARS-CoV-2 protein-coding gene. Host proteins that interacted with the expressed viral proteins within cells were identified by affinity purification followed by mass spectrometry (Gordon et al. 2020). The protein-RNA interaction network (Banerjee et al. 2020) was obtained from HEK293T cells individually transfected with SARS-CoV-2 expression constructs for each viral protein. Expressed viral proteins were cross-linked to their partner host RNA molecules and the protein-RNA complexes were selectively captured with the HaloLink Resin method. RNA-Seq libraries were prepared from the purified protein-RNA cross-link products and sequenced on an Illumina HiSeq 2500. Sequencing reads were then aligned to the human genome and genes whose RNAs were bound to viral proteins were identified.

### Quantification of viral reads

The NC_045512.2 release of the SARS-CoV-2 complete genome was downloaded from NCBI and the GRCh38.p13 human genome was obtained from the GENCODE website (Frankish et al. 2019). The raw reads of each RNA-Seq library were aligned with STAR (Dobin et al. 2013) to a reference genome composed of the human and SARS-CoV-2 genomes concatenated. The number of reads mapped to each genome in each library was calculated using an in-house script. The ratio of viral reads was calculated as the number of viral reads in each library divided by the total mapped reads in the library.

### Gene and transcript expression quantification

Transcript-level expression quantification was performed from the RNA-Seq raw reads using the alignment-independent software *salmon* (Patro et al. 2017) using the GENCODE GRCh38.p13 v33 human reference transcriptome (Frankish et al. 2019). The index was built with k-mers of size 31 and the quant function was run using default settings plus the *--validateMappings* and *--nBootstraps 100* flags. We also used the GENCODE v33 reference transcriptome (Frankish et al. 2019) file to create a transcript to gene (*tx2gene*) dictionary. This dictionary contained the correspondence between each transcript to its parent gene and the information of the biotypes of each gene and transcript. The package *tximport* was used to import *salmon* results to R and to summarize transcript-level to gene-level counts. Technical replicate sequencing libraries were collapsed by summing the counts of each transcript or gene from all replicates.

### Differential gene and transcript expression analysis

Gene and transcript-level count tables were used to perform differential expression analysis using DESeq2 (Love et al. 2014) with default settings. The design was set to be SARS-CoV-2 infected versus Mock samples in each experiment individually. The resulting genes and transcripts log_2_FoldChange values were shrunk with the *lfcShrink* function in DESeq2 using the *normal* mode (Love et al. 2014). Differentially expressed genes or transcripts were those with an adjusted p-value < 0.05 and abs(log_2_FoldChange) > 2. The *Isoformic* transcript-level analysis R pipeline was used to perform the experiments (https://github.com/luciorq/isoformic) (Queiroz et al. 2023).

### Sashimi plots

Sashimi plots were created to visualize splicing events in selected genes using the *ggsashimi* command-line tool (Garrido-Martín et al. 2018). Briefly, raw RNA-Seq reads were aligned to the GRCh38.p13 human genome using STAR (Dobin et al. 2013). The resulting BAM files and a GTF file of the same version of the genome were used to run *ggsashimi* for each selected gene. The genomic coordinates of selected genes were obtained using the *biomaRt* R package (Durinck et al. 2009).

### Custom gene set construction

We created custom sets of genes related to key biological processes and pathways involved in the innate immune response against viruses, antigen presentation via the class I MHC, the splicing machinery and regulation, the transport of RNA transcripts from the nucleus to the cytoplasm, and with NMD (Table S4). We built these custom gene sets using the terms and genes from the curated databases Reactome, Gene Ontology (biological processes, cellular compartment, and molecular function), MSigDB Hallmark pathways, KEGG, and Biocarta (Fig. S4). The authors selected the pathways and genes included in the custom gene set list based on their scientific experience.

### Functional enrichment analysis

Gene set enrichment analysis (GSEA) was performed for RNA-Seq results using the R package *fgsea (Korotkevich et al. 2016)* with the custom pathways created. For gene-level enrichment, we used the shrunk log_2_FoldChange values of all detected genes (no adjusted p-value cutoff) as recommended by the authors of the *fgsea* package (Korotkevich et al. 2016). Isoform-specific functional annotation databases similar to those built for genes are practically non-existing. Therefore, we extended the gene-level annotations of the selected genes in each custom pathway to their isoforms (Table S4) and ran separate GSEAs on *fgsea* with the shrunk log_2_FoldChange values of productive or unproductive transcripts separately. Significant pathways at the gene and transcript-levels were those with a p-value < 0.05.

DTPs were separated into up- and downregulated according to the log2FoldChange values provided by the authors who produced the data (Bojkova et al. 2020). Up- and downregulated DTPs were separately submitted to overrepresentation analysis (ORA) using the *fora* function in the *fgsea* package (Korotkevich et al. 2016). Significant pathways were those with a p-value < 0.05.

For the SARS-CoV-2 versus host protein-protein and protein-RNA interaction networks, we quantified the total number of host proteins or genes in each network that were members of the custom gene sets.

Functional enrichment and differential gene, transcript, and protein expression/translation results were represented using heatmaps, bar plots, and line plots created with *ggplot2* in R (Wickham 2016).

### Gene-level productive versus unproductive log2FoldChange plot

We built a gene-level productive versus unproductive log2FoldChange plot for all genes in the custom gene set. For each gene, productive or unproductive log2FoldChange values were calculated as the mean shrunk log2FoldChange value of all productive or unproductive isoforms of the gene. For example, the HLA-B gene has 8 annotated isoforms according to the GENCODE v33 reference transcriptome (Frankish et al. 2019): three protein-coding (productive) and 5 retained intron (unproductive). Thus, the productive fold change of the HLA-B gene was the mean log2FoldChange of its 3 productive isoforms, whereas the unproductive fold change of this gene corresponded to the mean log2FoldChange value of the 5 unproductive isoforms.

### Interaction networks

We used the *igraph (Csardi & Nepusz 2006)* and *ggraph* R packages to build and plot the unified virus versus host protein-protein network.

---

**Figure S1.**
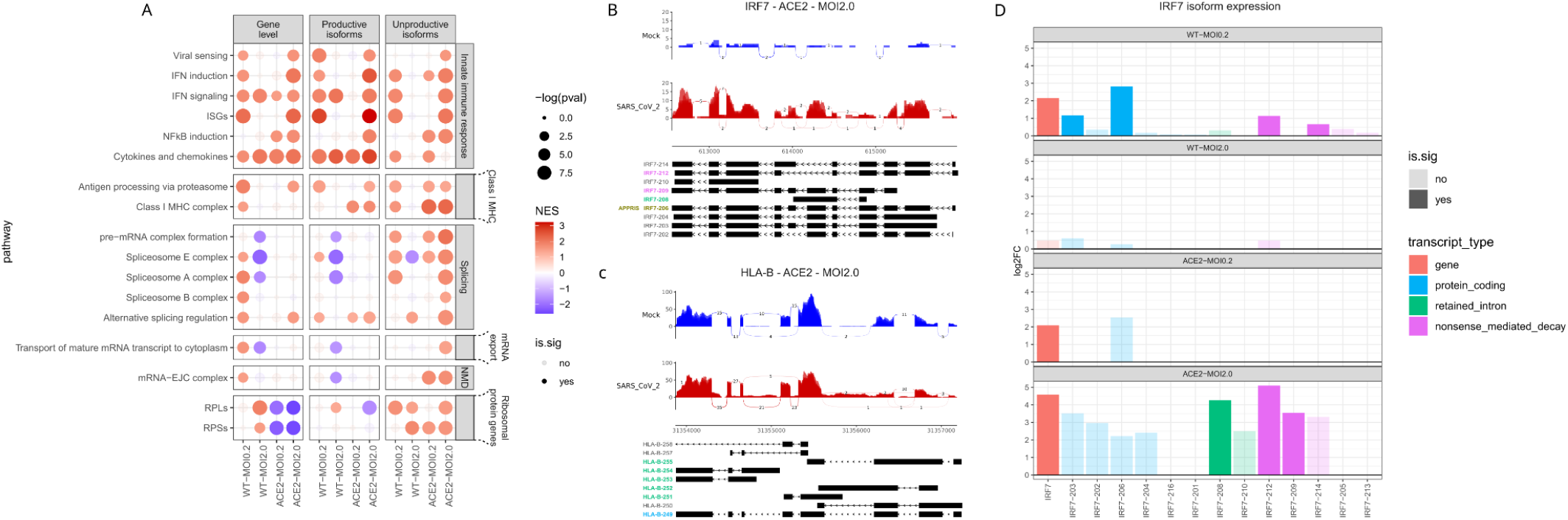
SARS-CoV-2 affects the expression of productive and unproductive transcripts in all cells. **A.** GSEA results for the DEGs (left panel) and productive and unproductive DETs (middle and right panels) in WT (MOI=0.2 and 2.0) and ACE2 (MOI=0.2 and 2.0) cells. The size of the dots is proportional to the -log_10_pvalue of the *fgsea* enrichment for each custom pathway and the color is related to the normalized enrichment score (NES - red positive, blue negative). **B, C.** Sashimi plots depicting RNA-Seq read coverage of *IRF7* (**B**) and *HLA-B* (**C**) genes in Mock (blue) and ACE2-MOI2.0 (red) cells. The structures of the transcripts of each gene that are expressed in ACE2-MOI2.0 are depicted under the sashimi plots. Transcript names are colored according to their types: green=retained intron, purple=NMD, gold=APPRIS main isoform, black=other non-DE transcripts. **D.** Gene (red) and transcript (blue=protein-coding, green=retained intron, and purple=NMD) log2FoldChange and significance status of IRF7 in WT (MOI=0.2 and 2.0) and ACE2 (MOI=0.2 and 2.0) cells.

## Notes

https://github.com/luciorq/isoformic

### Competing Interest Statement

The authors have declared no competing interest.

